# High-throughput phenotypic screen identifies a new family of potent anti-amoebic compounds

**DOI:** 10.1101/2021.10.06.463301

**Authors:** Conall Sauvey, Gretchen Ehrenkaufer, Jonathan Blevitt, Paul Jackson, Ruben Abagyan

## Abstract

*Entamoeba histolytica* is a disease-causing parasitic amoeba which affects an estimated 50 million people worldwide, particularly in socioeconomically vulnerable populations experiencing water sanitation issues. Infection with *E. histolytica* is referred to as amoebiasis, and can cause symptoms such as colitis, dysentery, and even death in extreme cases. Drugs exist that are capable of killing this parasite, but they are hampered by downsides such as significant adverse effects at therapeutic concentrations, issues with patient compliance, the need for additional drugs to kill the transmissible cyst stage, and potential development of resistance. Past screens of small and medium sized chemical libraries have yielded anti-amoebic candidates, thus rendering high-throughput screening a promising direction for new drug discovery in this area. In this study, we screened a curated 80,000-compound library from Janssen pharmaceuticals against *E. histolytica* trophozoites *in vitro*, and from it identified a highly potent new inhibitor compound. Further experimentation confirmed the activity of this compound, as well as that of several structurally related compounds, originating from both the Janssen Jump-stARter library, and from chemical vendors, thus highlighting a new structure-activity relationship (SAR). In addition, we confirmed that the compound inhibited *E. histolytica* survival as rapidly as the current standard of care and inhibited transmissible cysts of the related model organism *Entamoeba invadens*. Together these results constitute the discovery of a novel class of chemicals with favorable *in vitro* pharmacological properties which may lead to an improved therapy against this parasite and in all of its life stages.

**Author summary:** The parasite *Entamoeba histolytica* represents a significant challenge in the field of global health. It currently infects and causes disease among millions of people worldwide, particularly those lacking access to clean water. Drugs exist to treat this disease, but nevertheless it persists as a problem, likely at least partly due to problems and downsides inherent to these drugs. Hence the search for new and better ones is needed. We report here our contribution to this search, consisting of testing a large, carefully-curated collection of tens of thousands of chemicals for their ability to kill *E. histolytica*. This large-scale test resulted in the identification of one of the compounds as potently anti-amoebic, capable of killing the parasite cells at extremely low concentrations. Further experimentation found several chemically-related compounds to also possess this property, and additionally found the first compound capable of killing the infective life stage of another *Entamoeba* parasite. These results have revealed an entire new family of chemicals with good potential for development as better drugs against this disease.

## Introduction

*Entamoeba histolytica* is a parasitic protozoan amoeba that infects the human intestinal tract and causes the diarrheal disease amoebiasis, also known as amoebic colitis [1]. It is estimated to infect around 50 million people globally at any given time, resulting in approximately 50,000 to 70,000 deaths annually, and as such represents a significant problem from a global health perspective [2–4]. It exists in a two-stage life cycle consisting of an environmentally-resistant, infective cyst stage, and a mobile, invasive trophozoite stage. It is transmitted in a characteristic fecal-oral route, where cysts can be ingested from contaminated water or food. Once ingested, cysts pass through to the host intestinal tract where they release trophozoites. Trophozoites feed and multiply, and under certain conditions invade and infect the surrounding host tissues. They can further re-form into cysts which are passed in the host’s feces, and potentially onwards to other hosts [3]. Due to this mode of infection, amoebiasis is most widespread in places where fecal contamination of water or food is likely, such as those where water sanitation is insufficient or non-existent. Thus, it most heavily affects and damages populations that are already socioeconomically vulnerable [4–6]. Symptomatic amoebiasis occurs when trophozoites attack the intestinal lining, causing ulceration, and invade the surrounding tissues [7]. Symptoms usually include long-lasting diarrhea progressing to dysentery, as well as generalized abdominal tenderness and fever [1]. In extreme cases infection can spread from the intestinal region to others such as the liver, lungs, or brain [3, 8]. All of these are significantly more dangerous and more likely to result in mortality.

The current standard of care for amoebiasis is the nitroimidazole drug metronidazole, commonly known under the brand name “Flagyl.” Metronidazole has seen widespread use against both amoebiasis and other protozoan parasitic diseases since its discovery in the mid-20th century [9]. Despite its successes however, several critical issues exist with metronidazole which render necessary the continued search for new treatment options. One of these issues is the significant adverse effects that often accompany metronidazole treatment. Several of these adverse effects are similar to the symptoms of amoebiasis itself, such as diarrhea and fever, and can exacerbate the already difficult experience for the patient to the point of intolerability [10]. In fact, a study from a Rwandan group has found these effects to be associated with patient non-compliance with the course of metronidazole treatment, as well as worse clinical outcomes [11]. Another issue with metronidazole is its inability to kill the transmissible cyst stage of *E. histolytica*, necessitating followup treatment with an additional drug such as paromomycin in order to prevent disease spread [12]. This complication and extension of the overall course of treatment could possibly also reduce the likelihood of full patient compliance with the treatment regimen. Beyond these issues, a final concern is the possibility of emergent resistance to metronidazole therapy. While this has yet to be reported in the field, resistant strains of *E. histolytica* are routinely generated in laboratory settings [13–15].

Given these gaps in the effectiveness of metronidazole therapy for amoebiasis, the search for new and additional compounds capable of inhibiting this parasite is ongoing [16]. In particular, several screening efforts of various sizes have been undertaken, and have produced multiple compounds of interest [17, 18]. In (Debnath *et al*., 2012) the authors identified the FDA-approved drug auranofin as an inhibitor of *E. histolytica* trophozoite growth from a screen of the 910-member Iconix drug library. More recently, authors of this current study conducted a low-throughput targeted screen of antineoplastic kinase inhibitor drugs, identifying multiple highly-potent *E. histolytica* inhibitors including the cancer chemotherapy drug ponatinib [19]. Interestingly, another, much larger screen of the 11,948-member ReFRAME library conducted in parallel by authors of this study also identified ponatinib among its potent hits [18]. Given the success of these past screen-based studies, in this study we expanded upon their approach by conducting a high-throughput, semi-automated phenotypic screen of a large chemical library in collaboration with Janssen Pharmaceuticals, Inc. against *E. histolytica, in vitro*. This library, called the Jump-stARter library, consists of 81,664 small molecules selected for their favorable chemical properties for drug development purposes, as well as their structural diversity. The library was designed with these features in order to “jump-start” novel drug discovery efforts and allow for the identification of structures and structural features that are active in a specified context [20–25]. From the screen of this library, we found a highly active compound against *E. histolytica* trophozoites. Further investigation showed similar activity among several structurally-related compounds included in the library, as well as several additional such compounds purchased from commercial vendors. Of these compounds, we found nearly all of them to be non-toxic to a cultured human cell line. We also identified activity of the initial compound against cysts of the related model organism *Entamoeba invadens*, and found it to inhibit *E. histolytica* at the same rate as metronidazole. Together these results show this new compound to be a promising candidate for drug development against amoebiasis.

## Materials and methods

### Compound Library

The chemical compound library screened against *E. histolytica* trophozoites was obtained in collaboration with Janssen Pharmaceuticals. The library, referred to as the “Jump-stARter” library, contains a diverse collection of 80,000 drug-like small molecules intended for maximum potential efficacy in collaboration drug-discovery projects with external research groups [20–23, 25]. The Jump-stARter library was originally selected from millions of proprietary compounds by Janssen Pharmaceuticals medicinal chemists using “drug-likeness,” structural diversity, and favorable physical properties as criteria [24, 25]. For this study the library was spotted into black, clear-bottom 1536-well plates (Greiner) at 50nLper well in the Janssen compound logistics facility in Beerse, Belgium, and shipped directly to the University of California - San Diego.

### *Entamoeba* cell culture

*E. histolytica* strain HM-1:IMSS trophozoites were maintained in 50ml culture flasks (Greiner Bio-One) containing TYI-S-33 media, 10% heat-inactivated adult bovine serum (Sigma), 1% MEM Vitamin Solution (Gibco), supplemented with penicillin (100 U/mL) and streptomycin (100 μg/mL) (Omega Scientific) [17]. *E. invadens* strain IP-1 (a) were cultured in LYI-S-2 (b) at 25°C [26, 27].

### High-throughput phenotypic screen

Metronidazole and DMSO controls were added to the pre-spotted 1536-well Jump-stARter library plates using a Multidrop Combi reagent dispenser (Thermo Scientific), resulting in final assay concentrations of 25μM and 0.5% respectively. Cultured *E. histolytica* trophozoites were then seeded into the same plates at a density of 400 cells per well, also using a Multidrop Combi reagent dispenser, resulting in a final test compound concentration of 25μM. The plates were sealed into GasPak EZ (Becton-Dickinson) bags and incubated at 37°C for 48hr. 2μL of CellTiter-Glo luminescence-based cell viability assay reagent was added using a Multidrop Combi reagent dispenser, and plates were incubated in the dark for 10 minutes. Luminescence was measured using an EnVision plate reader (Perkin Elmer). Percent inhibition values for each well were calculated and plotted using Microsoft Excel. Outlying values and plates resulting from observed technical malfunctions such as clogging or bubbling during liquid handling were excluded from the final analysis.

### 384-well dose-response cell viability assay

Compounds at 5mM were transferred from a stock plate to black, clear-bottom 384-well plates (brand) and each diluted in DMSO in a 7-point series by a factor of 2. Diluted compounds were then transferred in triplicate into black, clear-bottom 384-well assay plates using an Acoustic Transfer System (EDC Biosystems). Final compound concentrations ranged from 25μM to 0.39μM. Cultured *E. histolytica* trophozoites were added at a density of 1000 cells per well, plates were sealed into GasPak EZ bags and incubated at 37°C for 48hr. CellTiter-Glo reagent was added and luminescence values were measured, and percent inhibition was calculated for each well as described previously.

### 96-well dose-response EC_50_ determination assay

Cultured *E. histolytica* trophozoites were seeded into white, solid-bottom 96-well plates (Greiner) at a density of 5000 cells per well. Test compounds were diluted in DMSO in 8-point series by a factor of 2 and added to wells in triplicate. 10μM metronidazole and 0.5% DMSO controls were added. Plates were sealed into GasPak EZ bags and incubated at 37°C for 48hr. CellTiter-Glo reagent was added and luminescence values were measured and percent inhibition was calculated for each well as described previously. EC_50_ values were calculated and dose-response curves were plotted using GraphPad Prism software.

### Human cell toxicity assay

HEK293 cells were cultivated in 75cm2 flasks (Corning) containing Dulbecco’s modified eagle medium (DMEM) (Gibco) supplemented with 10% fetal bovine serum (FBS) (Gibco) and antibiotic-antimycotic solution (Sigma-Aldritch). Cells were detached by trypsinization, collected, and seeded into (plates) at a density of 5000 cells per well. Test compounds were diluted in DMSO in a 6-point series by a factor of 2 and added to assay wells in quadruplicate alongside 0.5% DMSO and media-only controls. Final assay concentrations ranged from 25μM to 0.78μM. Assay plates were incubated for 48 hours, then CellTiter-glo reagent was added. Assay plates were incubated in the dark for 10 minutes, then luminescence values were read on an EnVision plate reader. Percent inhibition values for each replicate were calculated from luminescence data using Microsoft Excel, and LD50 values were calculated and dose-response curves plotted using GraphPad Prism.

### Timecourse assay

Cultured *E. histolytica* trophozoites were plated in 96-well plates at densities of 5000 cells per well. Compounds of interest were serially diluted by a factor of 2 in DMSO to a total of 8 points and added to wells in each replicate plate in triplicate along with 0.5% DMSO and 10μM metronidazole controls. The final in-well concentrations of the compounds of interest ranged from 25μM to 0.195μM. The replicate plates were individually incubated for either 6, 12, 24, 36, or 48 hours, following which CellTiter-glo reagent was added. Luminescence was measured, EC_50_ values were determined, and dose-response curves were plotted as described previously.

### Cyst viability assay

Mature cyst viability assay was performed as described previously, using a transgenic *E. invadens* line stably expressing luciferase (CK-luc) [16, 22]. Parasites were induced to encyst by incubation in encystation media (47% LG) [23]. After 72 h, parasites were washed once in distilled water and incubated at 25°C for 4–5 h in water to lyse trophozoites. Purified cysts were pelleted, counted to ensure equal cyst numbers, and resuspended in encystation media at a concentration of 1-5×105 cells per ml. One ml suspension per replicate was transferred to glass tubes containing encystation media and drug or DMSO, then incubated at 25°C for 72 h. On the day of the assay, cysts were pelleted and treated once more with distilled water for 5 h to lyse any trophozoites that had emerged during treatment. Purified cysts were then resuspended in 75 μl Cell Lysis buffer (Promega) and sonicated for 2×10 seconds to break the cyst wall. Luciferase assay was performed using the Promega luciferase assay kit according to the manufacturer’s instructions. Assays were performed on equal volume of lysate (35 μl) and not normalized to protein content. Effect of the drug was calculated by comparison to DMSO control, after subtraction of background signal. Significance of drug effects was calculated using a one-tailed T-test.

## Results

### High-throughput screen of the Jump-stARter library against *E. histolytica* trophozoites *in vitro*

In order to identify new inhibitors of the human parasite *E. histolytica* we established a collaboration with scientists from Janssen Pharmaceuticals, inc., and from them, obtained a copy of their Jump-stARter chemical library. This library consists of 81,664 small molecules selected by Janssen medicinal chemists for their chemical diversity and optimal physical properties for drug discovery and development efforts [20, 21, 24, 25]. We screened the Jump-stARter library against *E. histolytica* trophozoites in vitro using a semi-automated, high-throughput methodology. Previous screens using this organism have been accomplished using 96-well or 384-well plate formats, but due to the large size of the Jump-stARter library, we developed and utilized a novel 1536-well-plate-based methodology. All compounds in the library were tested at 25μM, and the viability of the parasite cells after incubation for 48 hours was measured with the luciferase-based CellTiter-glo assay. 297 compounds achieved greater than 70% inhibition values and were thus designated as ‘hits’ and selected for further investigation (Fig 1).

**Fig 1.**
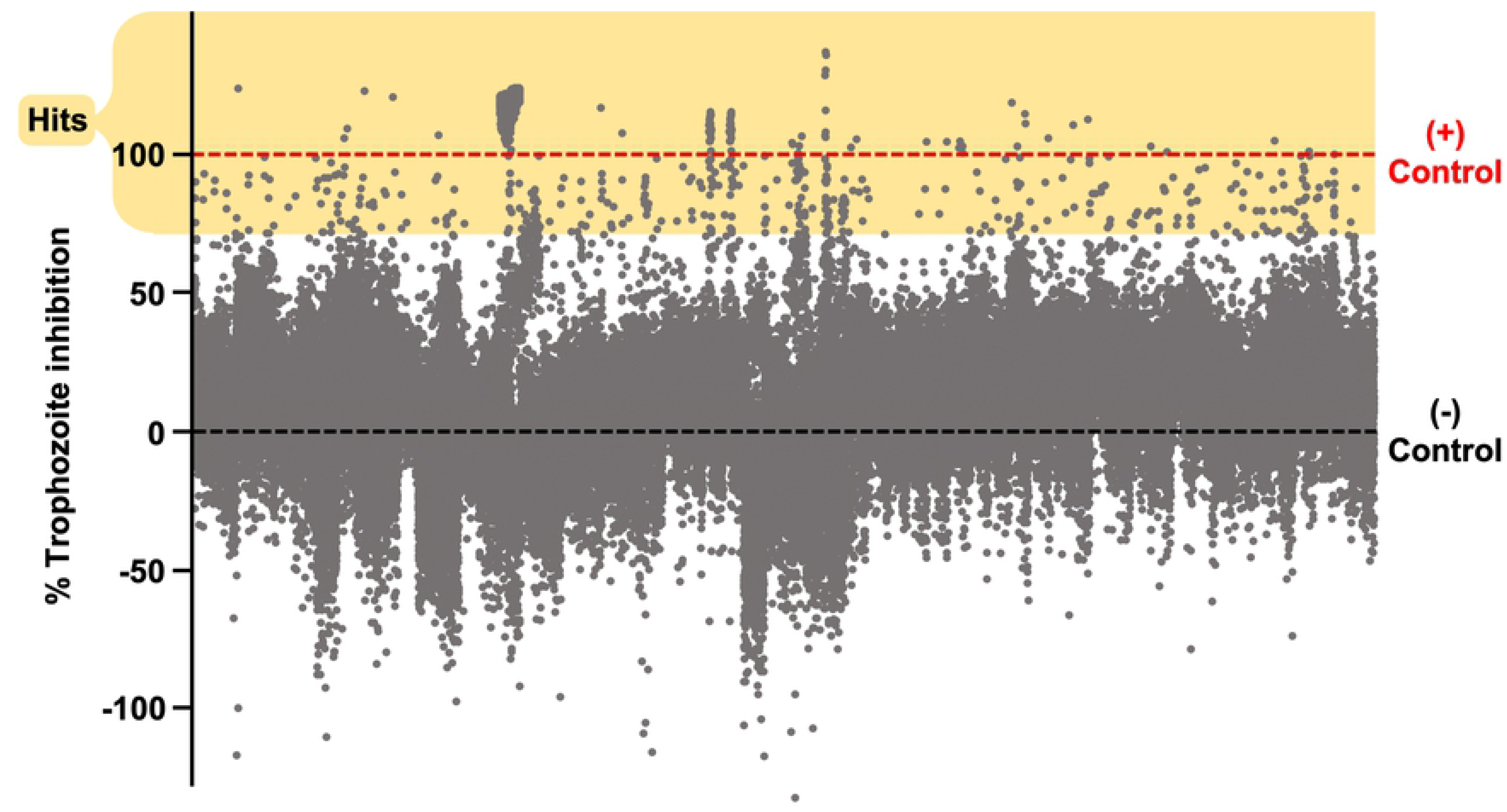
High-throughput screening results of 81,664 small molecules against *E. histolytica*. Scatterplot displays percent inhibition values of Jump-stARter library compounds against *E. histolytica* trophozoites calculated relative to positive and negative controls. Grey dots represent individual compound values. Dashed lines represent 0% (average of 10μM Metronidazole, positive controls) and 100% (average of 0.5% DMSO, negative control) percent inhibition values. Yellow box encloses compounds with a percent inhibition value greater than 70%, which were marked as hits and selected for further investigation.

### Dose-response screen of top candidate molecules to determine potency

Of the 297 hits from the high-throughput screen, 128 were available to be ordered in additional amounts from Janssen pharmaceuticals. In order to determine their potency, we ordered stock solutions of these compounds and conducted a medium-throughput dose-response screen against *E. histolytica* trophozoites *in vitro*. The compounds were diluted into triplicate 7-point dose-response curves ranging in final assay concentration from 25μM to 0.39μM in a 384-well plate format. Parasite cells were incubated with the compounds, and their viability measured using CellTiter-glo (S2 Dataset). 19 compounds which met the minimum criteria of either producing high levels of parasite inhibition across several points of concentration, or producing a varying range of inhibitions across the different concentrations were selected for further in-depth investigation. Compounds which achieved low or moderate levels of inhibition consistently across all points of concentration were excluded. In order to determine their EC_50_ values, the 19 compounds were tested against *E. histolytica* trophozoites in triplicate 8-point dose response curves in 96-well plates (Fig 2). From these, a compound designated hereafter as JNJ001 achieved the remarkably low EC_50_ value of 0.29μM, with the other compounds achieving values ranging from 4.30μM to 15.72μM (Table 1). These results revealed compound JNJ001 as the top hit, with high inhibitory potency against *E. histolytica* trophozoites.

**Table 1:** EC_50_ values of top hits from high-throughput screen. EC_50_ values determined from dose-response assay against *E. histolytica* trophozoites in vitro. Color scale indicates compound potency. Green = more potent, red = less potent, grey = no effect.

| Compound ID: | EC<sub>50</sub> (μM) |
| --- | --- |
| JNJ001 | 0.29 |
| JNJ002 | 4.30 |
| JNJ003 | 4.59 |
| JNJ004 | 4.59 |
| JNJ005 | 5.45 |
| JNJ006 | 5.78 |
| JNJ007 | 5.78 |
| JNJ008 | 6.09 |
| JNJ009 | 6.39 |
| JNJ010 | 8.37 |
| JNJ011 | 10.26 |
| JNJ012 | 10.92 |
| JNJ013 | 15.72 |
| JNJ014 | No effect |
| JNJ015 | No effect |
| JNJ016 | No effect |
| JNJ017 | No effect |
| JNJ018 | No effect |
| JNJ019 | No effect |

**Fig 2.**
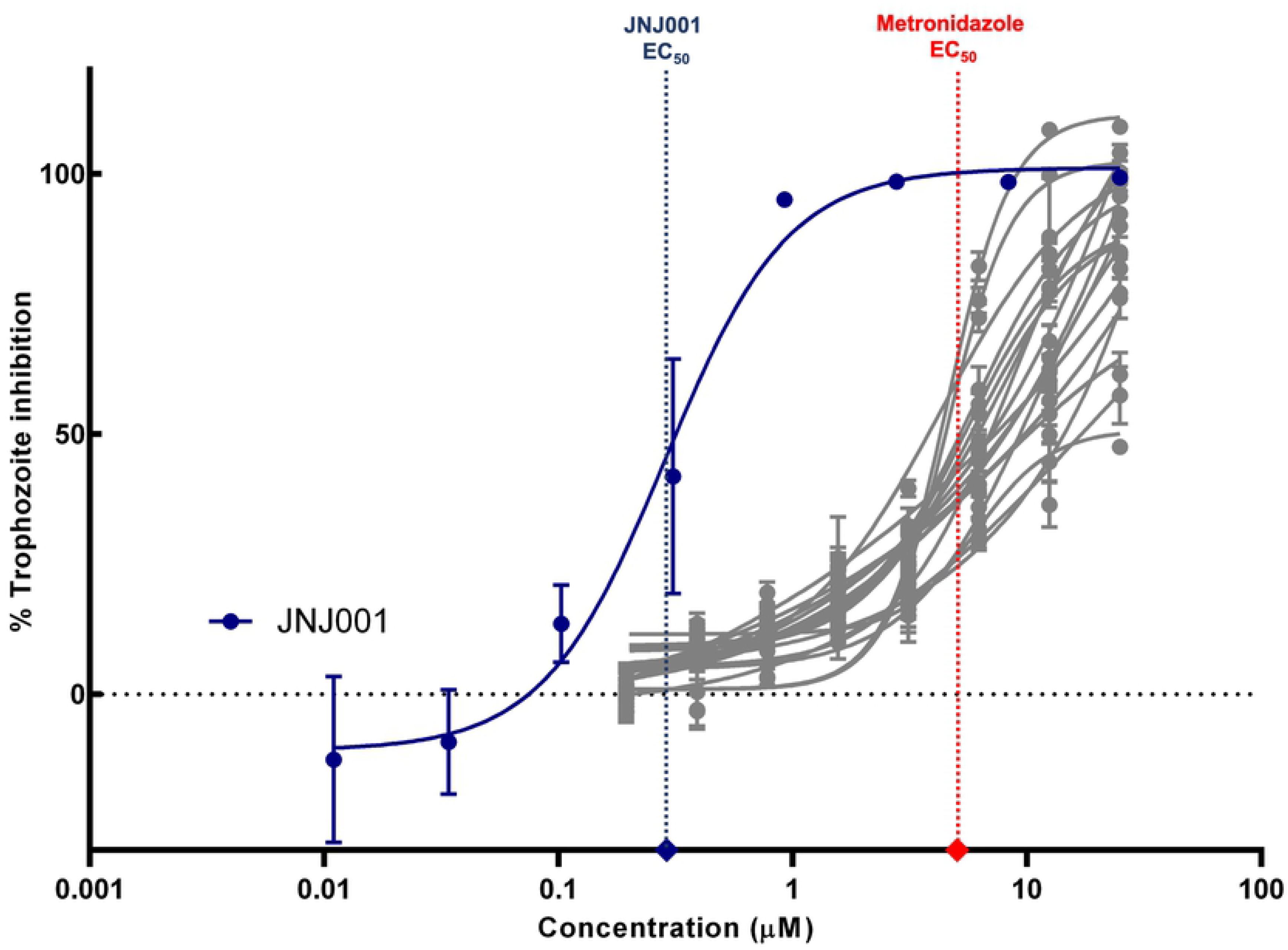
EC_50_ determination assay identifies a highly potent inhibitor of *E. histolytica*. Dose-response curves show percent inhibition of *E. histolytica* trophozoites compared to compound concentration. Trophozoite cell viability was measured after 48 hours of exposure to test compounds, and percent inhibition for each compound was calculated relative to controls. Compound JNJ001 (displayed in dark navy blue) was assayed over a broader range of concentrations compared to other compounds (displayed in grey) due to previous results indicating high potency in lower concentration ranges. Vertical dotted lines represent EC_50_ values of compound JNJ001 (navy blue) and metronidazole (red). Horizontal dotted line represents 0% inhibition level. Error bars represent standard deviation.

### Cell counting assay confirms inhibitory activity against *E. histolytica* trophozoites

In order to confirm the activity of compound JNJ001 against *E. histolytica*, it was once again tested in an 8-point dose-response curve in a 96-well plate, along with metronidazole as a control compound. In this assay however, survival of the parasites was directly determined by cell counting using a hemocytometer. When the results of this assay were plotted alongside the Luciferase-based dose-response curve results, they were found to match closely (Fig 3). This supports the accuracy of the luciferase-based assay results and indicates that compound JNJ001 does in fact act as a potent inhibitor of *E. histolytic*a trophozoites.

**Fig 3.**
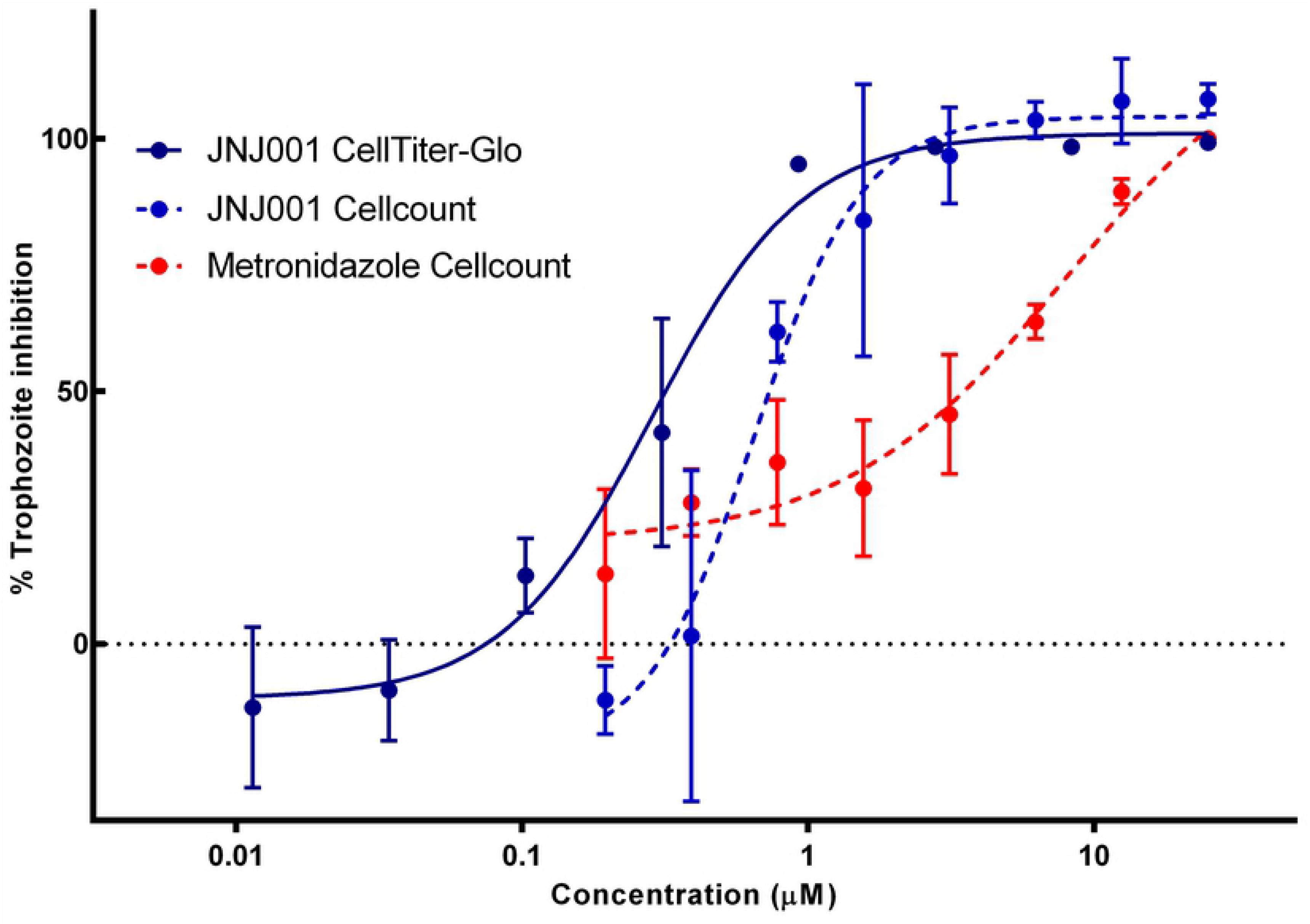
Cell counting assay confirms anti-amoebic activity of compound JNJ001. Dose-response curves show percent inhibition of *E. histolytica* trophozoites compared to compound concentration. Trophozoite cell viability or survival was measured after 48 hours of exposure to test compounds, and percent inhibition for each compound was calculated relative to controls. Dashed lines represent parasite survival measured by cell counting. Solid line represents parasite cell viability measured by a luminescence-based assay. Error bars represent standard deviation.

### Human cell toxicity assay

It is a crucial feature of any candidate compound in the drug development process that it not possess toxicity towards human cells, either specifically or as part of a general cytotoxicity. In order to determine whether compound JNJ001 possesses this feature, we tested it for toxicity against human HEK293 cells *in vitro*. In this experiment, human HEK293 cells were cultured and tested in a dose response assay with JNJ001, metronidazole and a selection of antineoplastic kinase inhibitor (AKI) drugs which have previously shown anti-amoebic activity [19]. Of these, only the AKI drug ponatinib produced notable toxicity to the human cells at concentrations up to 25μM, whereas JNJ001, the other AKI drugs, and metronidazole produced no toxicity at all (Fig 4). These results indicate that JNJ001 is both not specifically toxic to human cells and does not inhibit *E. histolytica* due to a generalized cytotoxic mechanism.

**Fig 4.**
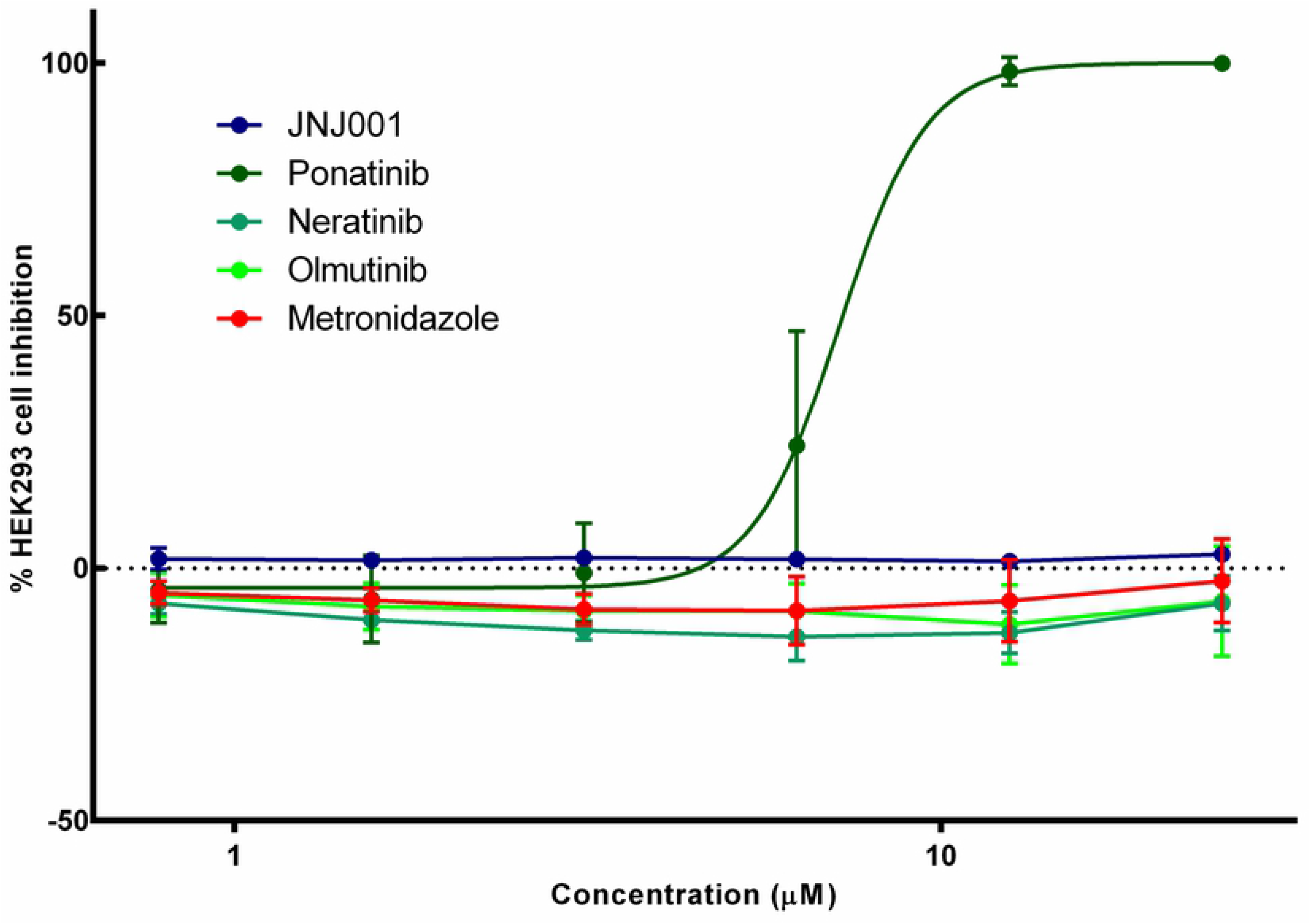
Compound JNJ001 does not inhibit human HEK293 cells. Dose-response curves show percent inhibition of human HEK293 cells compared to compound concentration. Cell viability was measured after 48 hours of exposure to test compounds, and percent inhibition for each compound was calculated relative to positive and negative controls. Compound JNJ001 is represented with a navy blue line. Metronidazole is represented with a red line. Ponatinib is represented with a green line. Error bars represent standard deviation.

### Compound JNJ001 inhibits *E. histolytica* as rapidly as metronidazole

In order to determine the speed with which compound JNJ001 achieves its inhibitory potency against *E. histolytica*, we measured and calculated its EC_50_ values at a series of timepoints subsequent to administration, alongside metronidazole for comparison. Cell viability was measured using CellTiter-glo at 12, 24, 36, and 48 hours after the addition of compounds. Both JNJ001 and metronidazole achieved their lowest EC_50_ values between the 24 and 36 hour timepoints (Fig 5), indicating that JNJ001 acts to inhibit *E. histolytica* as rapidly as the current standard of care drug.

**Fig 5.**
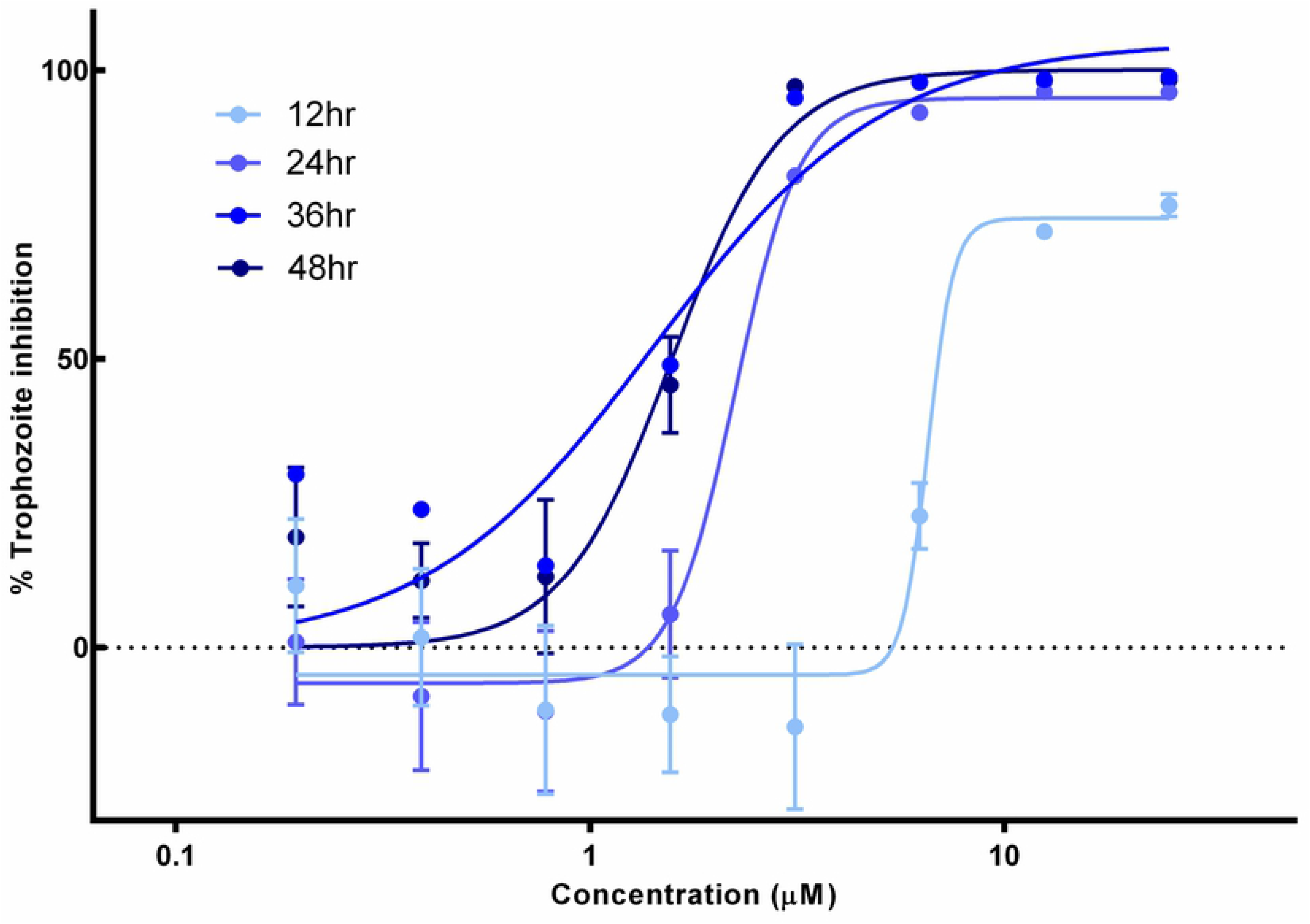
Compound JNJ001 achieves peak anti-amoebic effects within 24-36 hours. Dose-response curves show percent inhibition of *E. histolytica* trophozoites compared to compound concentration. Trophozoite cell viability or survival was measured after 12, 24, 36, and 48 hours of exposure to test compounds. Percent inhibition for each data point was calculated relative to average positive and negative control values. Error bars represent standard deviation.

### Compound inhibits mature *Entamoeba* cysts

As *E. histolytica* cannot be induced to encyst in vitro, we utilized the related parasite, *E. invadens*, a well-characterized model system for *Entamoeba* development, to assay for inhibition by compound JNJ001 [28]. Mature (72h) cysts of a transgenic line constitutively expressing luciferase were treated with JNJ001 at 10μM or 0.5% DMSO as negative control, for 3 days. After treatment, cysts were treated with distilled water for five hours to remove any remaining trophozoites, and luciferase activity was assayed. JNJ001 was found to reduce luciferase signal compared to controls, indicating that it is capable of killing *Entamoeba* cysts (Fig 6). In contrast, metronidazole up to 20 μM has been previously demonstrated to produce no such inhibition [19].

**Fig 6.**
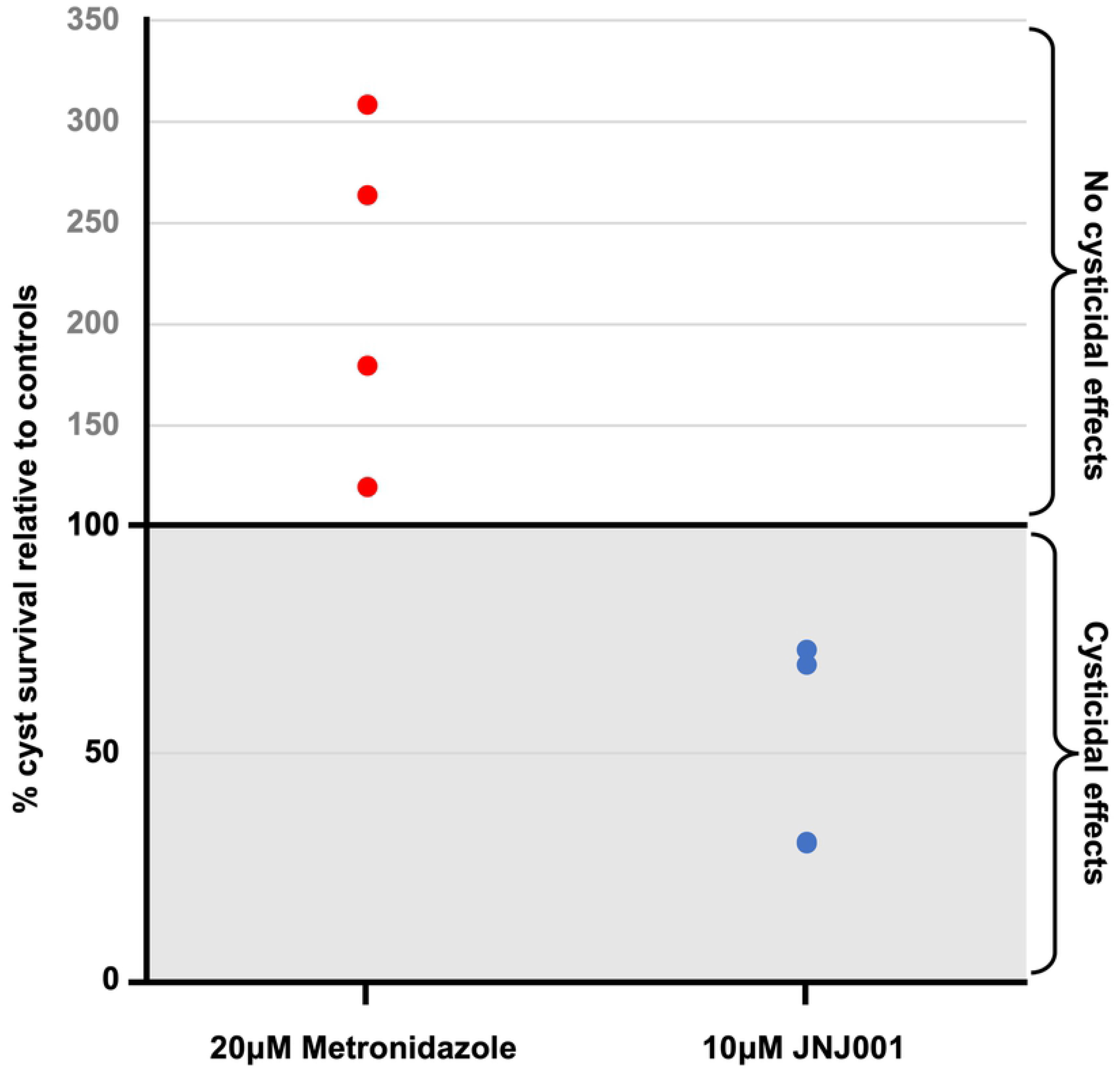
Compound JNJ001 inhibits *E. invadens* cysts. Graph representing survival of luciferase-expressing *E. invadens* cysts treated with 10μM compound JNJ001 (navy blue markers) or 20μM metronidazole (red markers) relative to 0.5% DMSO-treated controls. Markers represent individual luciferase readings. Metronidazole data taken from (Sauvey *et al*., 2021) [19].

### Screen of compounds structurally related to JNJ001 against *E. histolytica* trophozoites

In order to find additional inhibitors of *E. histolytica* based on compound JNJ001, we searched for structurally related compounds in both the hit list from the Jump-stARter library and online chemical vendor catalogs. We began by conducting a structural clustering analysis of the 128 available Jump-stARter library hits from the high-throughput screen. From this clustering analysis we identified 9 structurally similar series expansion compounds (designated as JNJ001-SE01 to JNJ001-SE09) and determined their EC_50_ values using *in vitro* dose-response assays as previously described (Figs 7,8). The resulting values ranged 1.38μM to 6.57μM (Table 2). We next conducted substructure- and similarity-based searches of online chemical vendor catalogs and identified 14 purchasable compounds (designated as CAB01-CAB14) similar to JNJ001. These were then purchased and tested to determine their EC_50_ values (Figs 9,10). The resulting values ranged from 4.53μM to 44.85μM (Table 3). These results together indicate that compound JNJ001 is a member of a structurally-related family of small molecules with anti-amoebic properties.

**Table 2.**
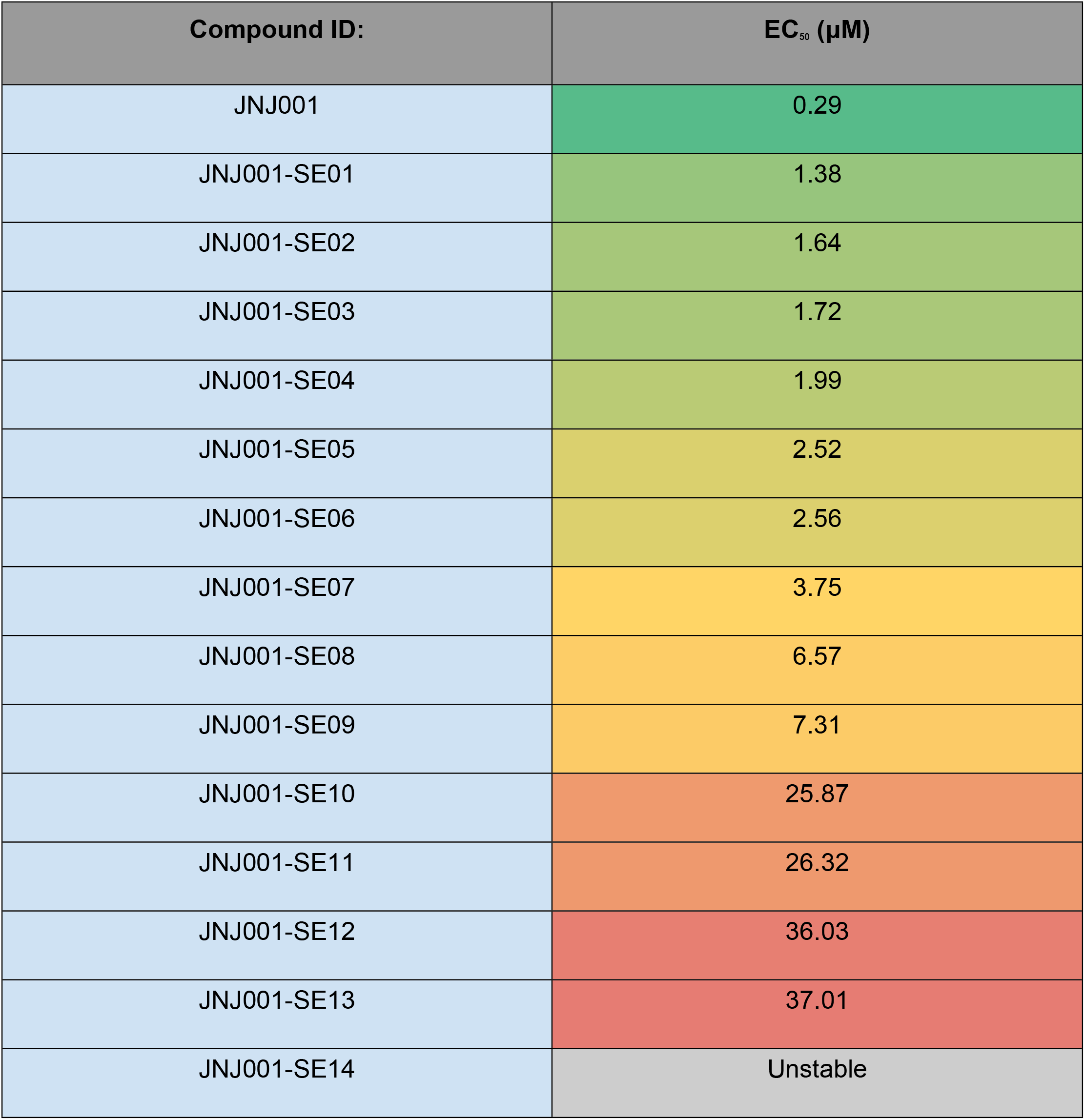
EC_50_ values of Jump-stARter library JNJ001 series expansion compounds against *E. histolytica* trophozoites in vitro. EC_50_ values determined from dose-response assay against *E. histolytica* trophozoites in vitro. Color scale indicates compound potency. Green = more potent, red = less potent, grey = no effect.

**Table 3.** EC_50_ values of non-Jump-stARter library JNJ001 series expansion compounds against *E. histolytica* trophozoites in vitro. EC_50_ values determined from dose-response assay against *E. histolytica* trophozoites in vitro. Color scale indicates compound potency. Green = more potent, red = less potent, grey = no effect.

| Compound ID: | EC<sub>50</sub> (μM) |
| --- | --- |
| CAB07 | 4.53 |
| CAB10 | 6.27 |
| CAB09 | 7.40 |
| CAB05 | 8.33 |
| CAB11 | 10.81 |
| CAB12 | 12.46 |
| CAB02 | 12.91 |
| CAB13 | 14.32 |
| CAB14 | 15.34 |
| CAB08 | 17.34 |
| CAB04 | 22.86 |
| CAB06 | 23.39 |
| CAB01 | 24.17 |
| CAB03 | 44.85 |

**Fig 7.**
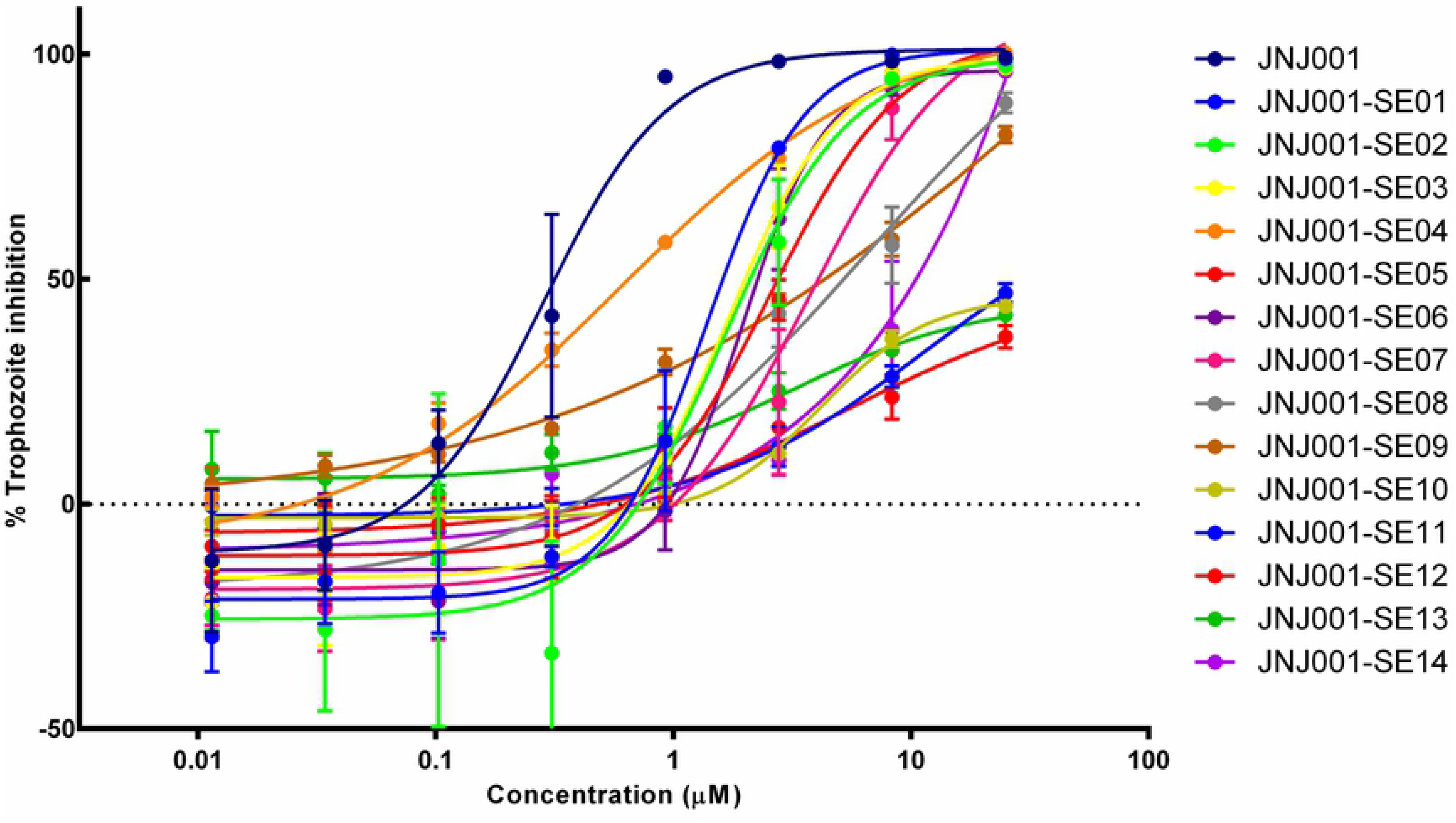
Dose-response assay results for Jump-stARter library JNJ001 series expansion compounds. Dose-response curves show percent inhibition of *E. histolytica* trophozoites compared to compound concentration. Trophozoite cell viability or survival was measured after 48 hours of exposure to test compounds, and percent inhibition for each compound was calculated relative to controls. Compound JNJ001 data (shown as a navy blue line) from Figure 2 is included for comparison. Error bars represent standard deviation.

**Fig 8.**
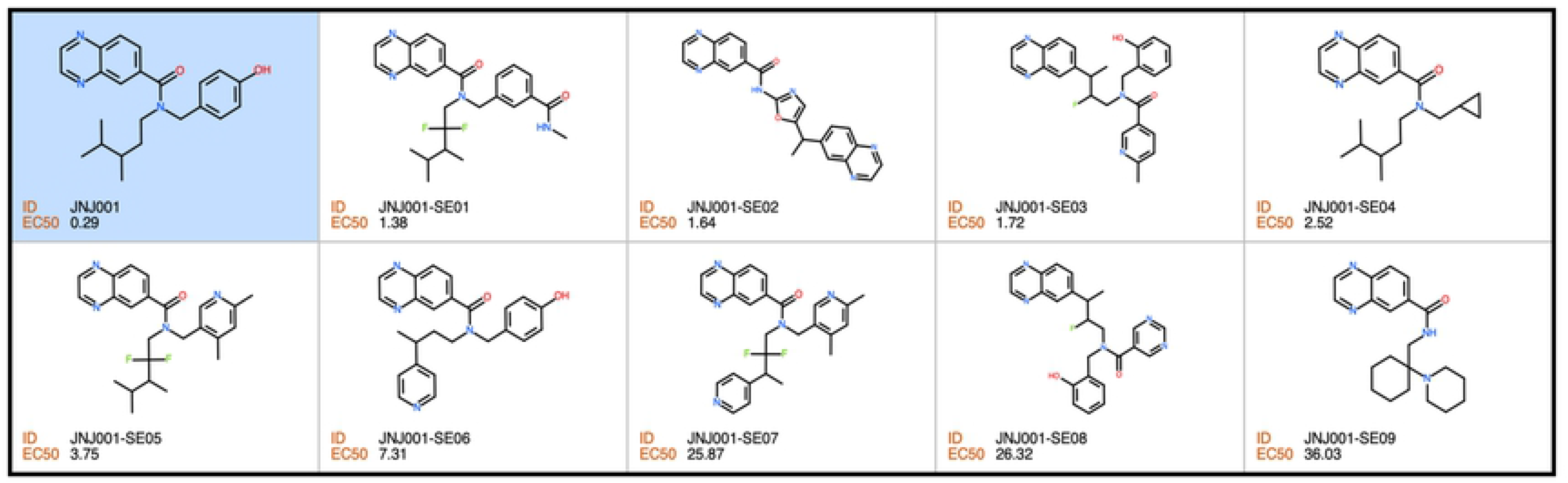
Structures of Jump-stARter library JNJ001 series expansion compounds. Structures, IDs, and measured EC_50_ values of compounds structurally related to JNJ001 (highlighted in blue) found in the Janssen Jump-stARter library.

**Fig 9.**
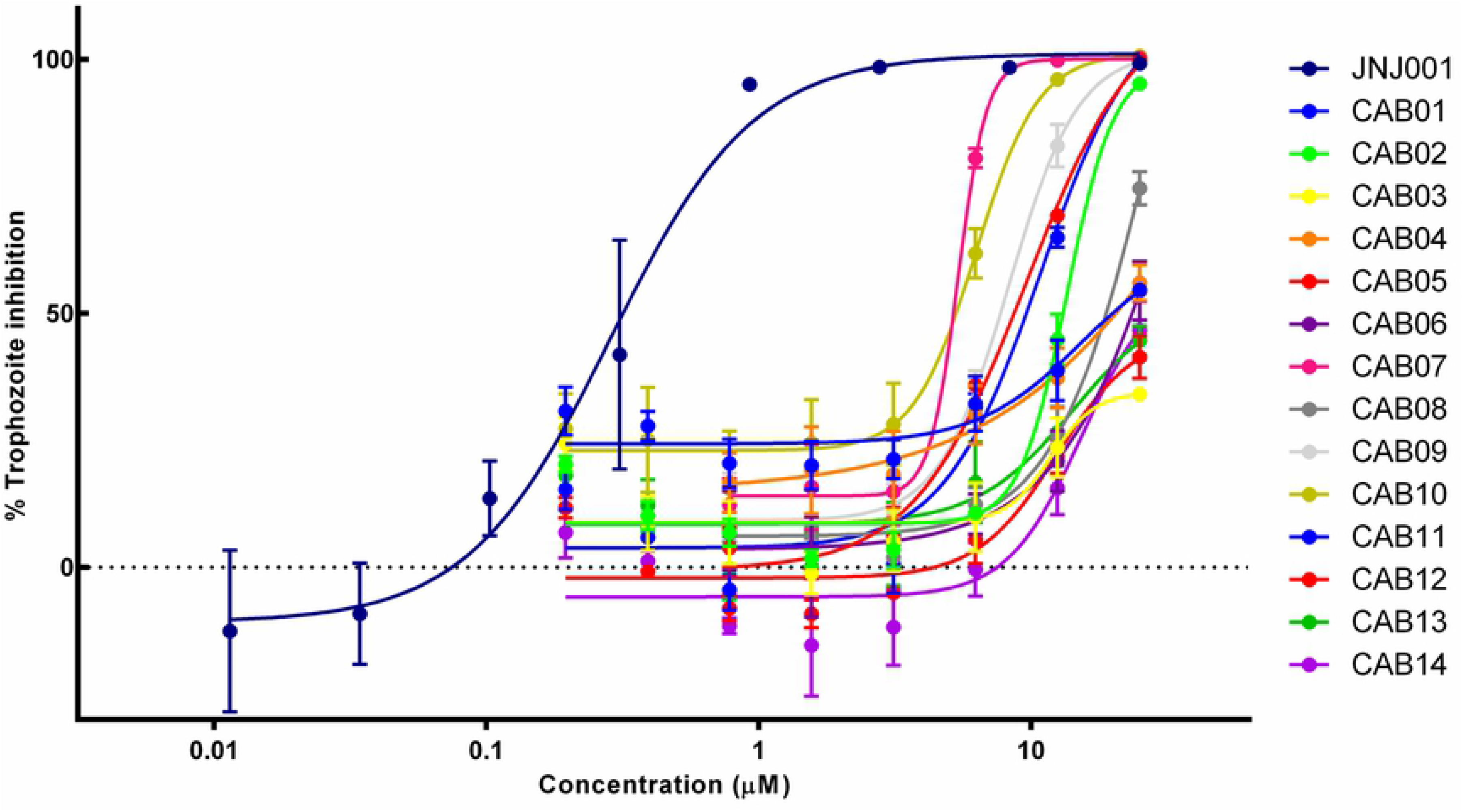
Dose-response assay results for non-Jump-stARter library JNJ001 series expansion compounds. Dose-response curves show percent inhibition of *E. histolytica* trophozoites compared to compound concentration. Trophozoite cell viability or survival was measured after 48 hours of exposure to test compounds, and percent inhibition for each compound was calculated relative to controls. Compound JNJ001 data (shown as a navy blue line) from Figure 2 is included for comparison. Error bars represent standard deviation.

**Fig 10.**
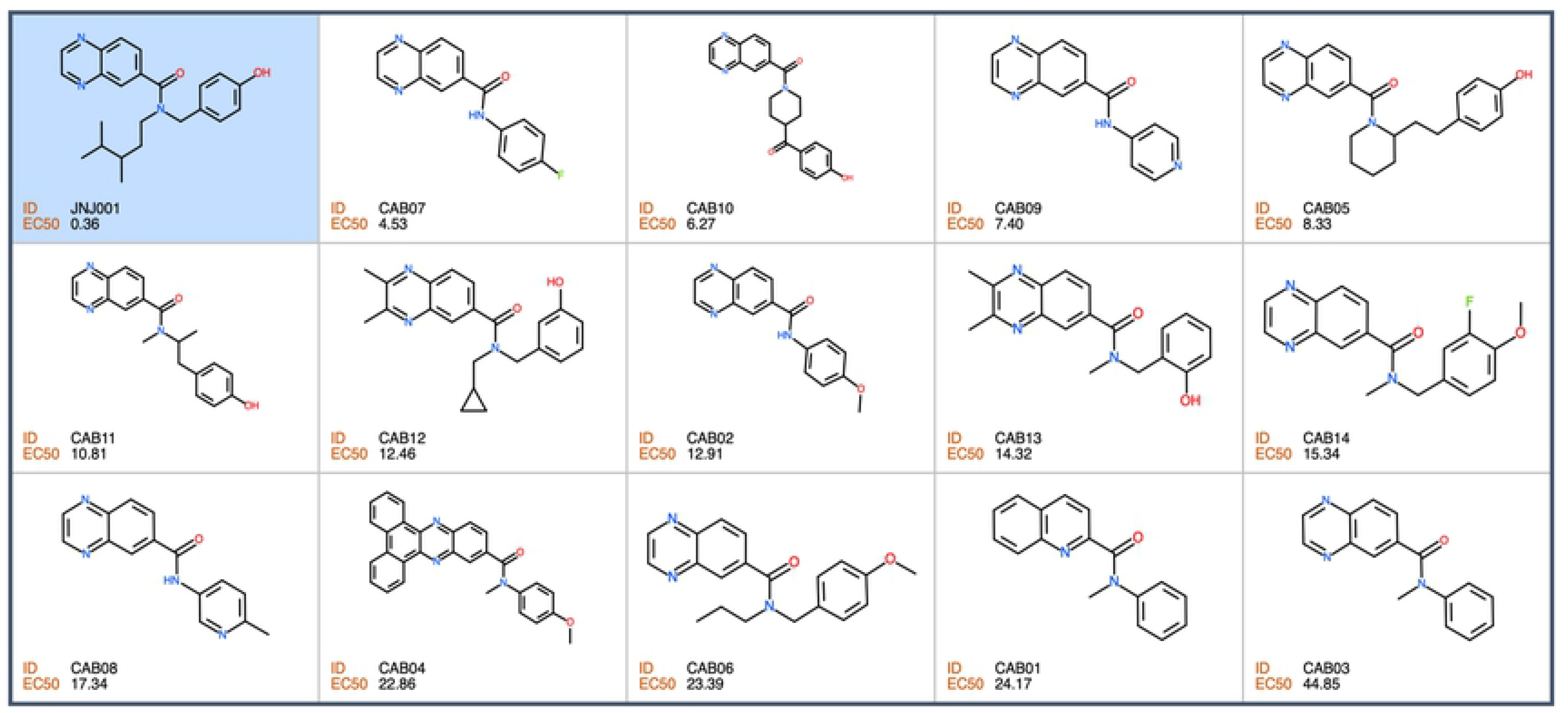
Structures of non-Jump-stARter library JNJ001 series expansion compounds. Structures, IDs, and measured EC_50_ values of additional compounds structurally related to JNJ001 (highlighted in blue).

### Human cell toxicity assay of series expansion compounds

In order to determine whether, like JNJ001, the series expansion compounds are cytotoxic to human cells, we tested them in a dose-response assay against cultured human HEK293 cells. Of the compounds tested, none produced notable toxicity to the human cells at concentrations up to 25μM (Fig 11). These results indicate that the compounds structurally-related to JNJ001 are not inhibiting *E. histolytica* due to a generalized cytotoxicity.

**Figure 11.**
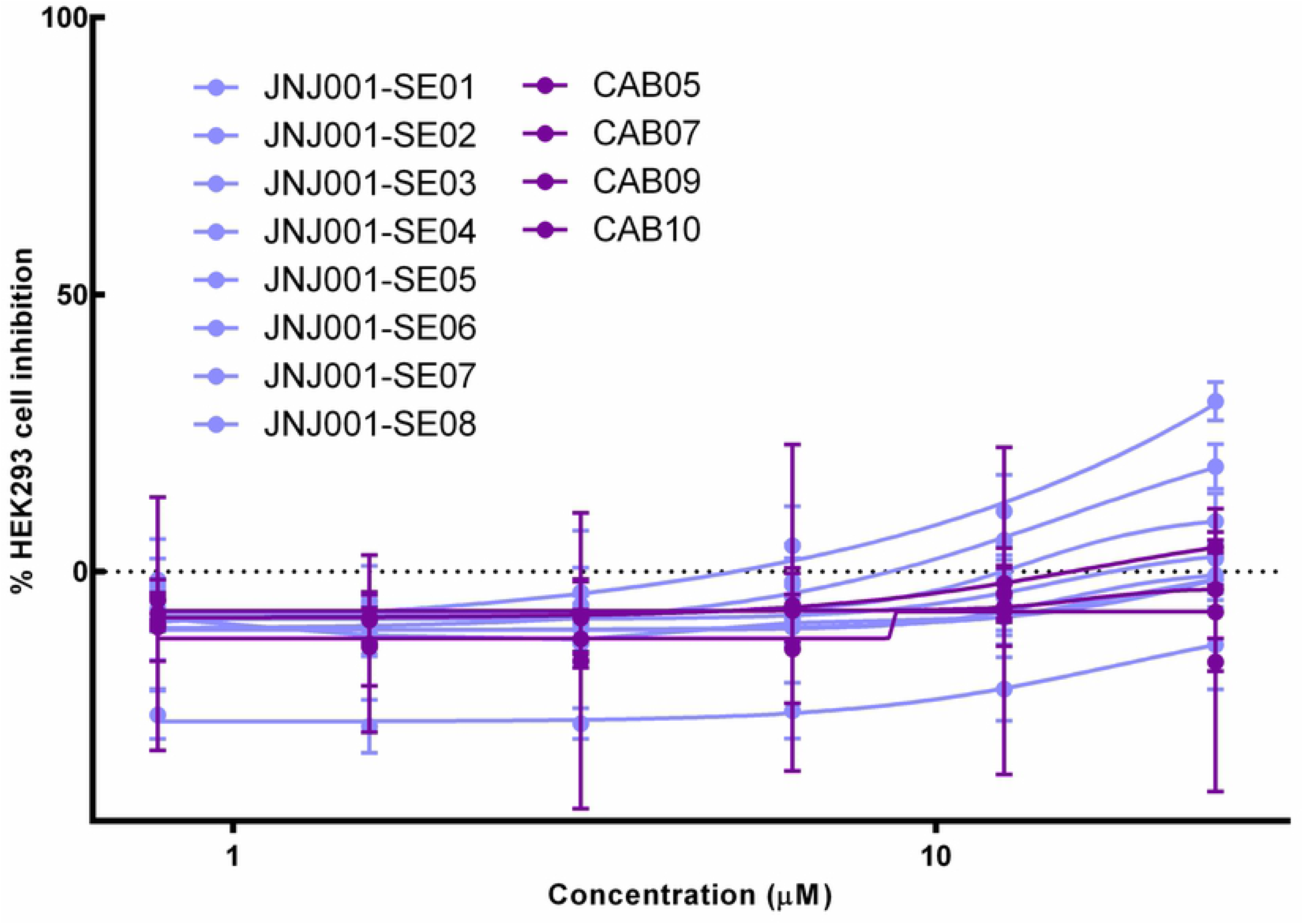
Series expansion compounds do not cause cell death in human HEK293 cells. Dose-response curves show percent inhibition of human HEK293 cells compared to compound concentration. Cell viability was measured after 48 hours of exposure to test compounds, and percent inhibition for each compound was calculated relative to positive and negative controls. Jump-stARter library compounds structurally related to JNJ001 are represented with light blue lines. Purchased compounds structurally related to JNJ001 are represented with pink lines. Error bars represent standard deviation.

## Discussion

Amoebiasis caused by infection with *E. histolytica* remains a significant public health issue in many places throughout the world, despite the longstanding availability of the drug metronidazole as a treatment [3, 4]. Multiple factors may contribute to this, including infrastructural problems in affected areas such as difficulties with water and food sanitation [4]. However, another important factor may be the shortcomings of metronidazole itself. Strong side effects and metronidazole’s inability to kill transmissible cysts complicate the course of treatment and have been suggested to contribute to patient non-compliance and resulting increased disease spread [10, 11]. As such, research into new and better anti-amoebic drugs is ongoing. For much of the history of this research, efforts have focused on inhibiting or disrupting specific cellular processes or targets within the parasite such as redox metabolism or kinase proteins [16]. As an example, authors of this study identified several antineoplastic kinase inhibitor drugs as highly potent inhibitors of *E. histolytica* [19]. More recently however, some groups, including authors of this study, have begun to achieve success with non-targeted screens ranging from low-to-high-throughput [17, 18]. In the current study, we continued and expanded upon this approach by conducting a large high-throughput in vitro screen against *E. histolytica*, resulting in the identification of a promising new family of potent inhibitors of this parasite.

The primary objective of this study was to use a high-throughput screening approach to identify novel small-molecule inhibitors of *E. histolytica* with high potential for further development as treatments for amoebiasis. To achieve this, we began by establishing a set of specific properties which would define an ideal successful candidate compound in our screen. We determined that such a compound would need to meet the following criteria: Firstly, to possess “drug-like” chemical properties such as appropriate molecular weight, solubility, and lack of known toxic groups. Such properties would be necessary for any further research efforts seeking to develop the compound into a useful drug treatment for amoebiasis. Secondly, to surpass the current standard of care drug metronidazole in potency. A higher potency could help address several of the drawbacks of metronidazole, such as lowering the dosage required, and potentially reducing adverse effects as a result.

Thirdly, to equal or surpass metronidazole in other measurable attributes such as rapidity of action and non-toxicity to human cells. These are areas that metronidazole performs well in, and any candidate compound would need to at least match it in order to be considered further. Fourthly, to exhibit inhibitory activity towards *Entamoeba* cysts, unlike metronidazole. A compound capable of any such inhibition would represent a large advantage over metronidazole, as metronidazole’s inability to kill cysts is currently one of its primary drawbacks.

With these four criteria established, we next proceeded to design the overall experimental approach with them in mind. We addressed the first criteria in the core of our design by selecting the Jump-stARter chemical library as the source of compounds for screening. This library was compiled by scientists at Janssen Pharmaceuticals specifically for use in novel drug discovery efforts and has been used in multiple collaborative projects of that nature [22, 23]. The compounds in the library were selected for their “drug-likeness,” having desirable chemical properties for drug development research closely matching those described in our first criteria. They were also selected for structural diversity and clustering into groups of related compounds, thus facilitating expansion of a single hit into a series or family of molecules. The library is also the largest yet screened against *E. histolytica* at over 80,000 compounds, which further increased the probability of identifying a high-potency hit, and thus helped address the second criteria.

However, this large library size also represented the next important challenge in our experimental design. Previous screening efforts against *E. histolytica* have used a maximum number of wells per assay plate of 96 or 384 [17, 18]. In the case of a library of this size, using plates of those well densities would necessitate using hundreds of them, which would result in a lengthy and complicated screening procedure. In order to avoid this and streamline our process we developed and optimized a screening procedure using 1536-well plates, thus drastically reducing the overall number of plates required, and hence the time required for screening them. This technical advance also has the additional benefit of opening up new possibilities of scale for future screening efforts against *E histolytica*.

Once the assay protocol was established, we screened the Jump-stARter library against *E. histolytica* trophozoites *in vitro* using a luciferase-based cell viability assay, which effectively measures the adenosine triphosphate (ATP) contained within surviving cells in each reaction volume, which can in turn be compared with average positive and negative control values to obtain a percent inhibition for each compound screened. We then further tested top-scoring compounds in assays to determine their EC_50_ values, revealing one in particular, compound JNJ001, as highly potent. We calculated the EC_50_ value of this compound as 0.29μM - more than a full order of magnitude greater than the previously established values for metronidazole (2-5 μM), and hence fulfilling our second desired criteria [19]. We confirmed this result with a secondary assay using a direct cell-counting method to show the compound as a true positive hit independent of assay type.

Having identified compound JNJ001 as fulfilling the first two of our desired criteria, we assayed the compound for toxicity to human HEK293 cells in vitro. We found compound JNJ001 (similarly to metronidazole) to be completely non-toxic at doses up to 25μM, and far exceeding its EC_50_ value. This effectively resulted in a wide therapeutic range spanning from the low tens of nanomolar into the tens of micromolar. We also performed a timecourse assay to compare the compound’s rapidity of action with that of metronidazole, and found both it and metronidazole to achieve their final EC_50_ values within 24-36 hours. Together these results confirmed that compound JNJ001 satisfied our third criteria.

Next, we approached our fourth criteria by assaying the ability of compound JNJ001 to inhibit the cysts of *Entamoeba invadens*, a closely-related reptilian parasite. In our experiment, compound JNJ001 was found to moderately inhibit *E. invadens* cysts, whereas metronidazole did not at all. This moderate cysticidal activity represents a significant advantage over metronidazole.

Having successfully identified a compound which fulfilled all of the desired criteria, we next sought to expand upon these results and search for additional related molecules with anti-amoebic activity. We used structural clustering to identify several related molecules within the Jump-stARter library. We also used substructure searching to find and obtain additional structurally-related molecules from commercial sources. We then took these two sets of series expansion compounds and assayed them for both their EC_50_ against *E. histolytica* trophozoites, and their toxicity towards cultured human cells. We found several of the compounds to possess good EC_50_ values, though not as potent as compound JNJ001, and additionally found all of them to possess very low toxicity to human HEK293 cells. These results served as further validation of the anti-amoebic activity of compound JNJ001, confirming that it was not just an isolated phenomenon, but instead can be attributed to a cluster of related molecules with shared activity against this parasite. This discovery also greatly broadened and diversified our set of available anti-amoebic compounds, providing several additional options for further development, some of which may prove to possess even more desirable chemical properties for that process.

Taken together the results described in this study indicate the existence of a promising new family of safe, structurally-related small molecules with strong anti-amoebic properties against both trophozoites and cysts, and good potential for further development as amoebiasis drugs. Going forward, several areas of further research are now possible. Further studies on the efficacy of these compounds in animal models of *E. histolytica* infection would be valuable, though this will be rendered challenging by the difficulties of establishing and maintaining such models [29, 30]. Similarly, safety and tolerability studies in cultured cells, animal models, and humans would be an essential aspect of developing these into clinical treatments. The possibility also exists and can be explored that these compounds may be effective against other related parasitic species. In conclusion, the discovery of these new anti-amoebic compounds represents an exciting new opportunity in the area of amoebiasis research.

## Acknowledgments

The authors would like to thank the following for their contributions to this work:

Monica Mendes Kangussu Marcolino for her assistance with *Entamoeba invadens* culture and screening

Kirti Khandwal Chahal for providing human HEK293 cells

Jean Bernatchez and Danielle Skinner for their assistance with screening equipment training Jim McKerrow for establishing and maintaining the high-throughput screening core and equipment as head of the Center for Discovery and Innovation in Parasitic Diseases at the Skaggs School of Pharmacy and Pharmaceutical Sciences at the University of California – San Diego

---

**S1 Dataset. Fig1 data.**

**S2 Dataset. 384-well dose-response screen data.**

**S3 Dataset. Fig2 data.**

**S4 Dataset. Fig3 data.**

**S5 Dataset. Fig4 data.**

**S6 Dataset. Fig5 data.**

**S7 Dataset. Fig6 data.**

**S8 Dataset. Fig7 data.**

**S9 Dataset. Fig9 data.**

**S10 Dataset. Fig11 data.**

## Notes

### Competing Interest Statement

The authors have declared no competing interest.

